# The impact of sex and ancestry on the molecular landscape of pancreatic adenocarcinoma

**DOI:** 10.64898/2026.10.06.757201

**Authors:** Caroline Y. Chen, Jaron Arbet, Chenghao Zhu, Jieun Oh, Paul C. Boutros

## Abstract

**Background:** Pancreatic ductal adenocarcinoma (PDAC) is projected to become the second leading cause of cancer-related mortality, with an increasing incidence and dismal 10% survival at 5 years despite best available care. PDAC is characterized by low tumour cellularity, recurrent driver mutation alterations, and substantial inter and intra-tumoural heterogeneity. The influence of individual genetic features such as sex and ancestry on tumour heterogeneity remains poorly understood.

**Methods:** We assembled a cohort of PDAC across five datasets for a total of 7,527 tumours. Measures of genomic instability were assessed across sex and ancestry on a global level (SNV density and percent genome altered by copy number events), and at a gene level for recurrent events. KRAS variant frequencies, ploidy and mRNA abundance were also evaluated. Ancestry was assigned using ADMIXTURE from germline genotypes where available, and patient reported otherwise.

**Results:** We observed ancestry-specific mutation patterns in global point mutation density, with higher SNV burden in African PDAC tumours and lower SNV burden in Asian PDAC tumours as compared to European tumours. We also identified an ancestry bias in the driver gene, *GNAS*, in which driver SNVs in *GNAS* was detected more frequently in those of Asian ancestry, and less frequently in those of African ancestry. No sex- or ancestry-associations were found for *KRAS*, but individual *KRAS* variants showed a trend towards ancestry-related variation.

**Conclusions:** These findings demonstrate that ancestry contributes to the genomic heterogeneity of PDAC, shaping both global and gene-specific mutation patterns. This work underscores the importance of including diverse ancestries in PDAC molecular profiling efforts which may inform the development of tailored molecular diagnostics and targeted therapies.

## Background

Pancreatic ductal adenocarcinoma (PDAC) is a lethal cancer of the exocrine pancreas with ∼90% five-year mortality and is projected to become the second leading cause of cancer death by 2030. Due to its rapid evolution, non-specific presenting symptoms and ineffectual screening, 80-85% of cases are unresectable or metastatic at presentation [1,2]. PDAC is characterized by low tumour cellularity, highly recurrent driver mutations and substantial inter- and intra-tumoural heterogeneity beyond these driver mutations [3]. The causes of inter-tumoural heterogeneity and their contributions to clinical outcomes remains unclear.

In other cancers, significant portions of inter-tumoural heterogeneity can be rationalized by considering patient characteristics, particularly germline variation. For example in prostate cancer, somatic *TMPRSS2*-*ERG* fusions are less frequent in patients of African ancestry [4]. PDAC has a significant germline component: ∼10% of patients carry a deleterious germline variant, most commonly *BRCA2* (∼3%), *ATM* (∼2%) or *BRCA1* (∼1%) [5]. We quantified the effects of the two largest genetic variations – sex and ancestry – on the inter-tumoural heterogeneity and mutational landscape of PDAC.

## Methods

### Experimental design

This study was designed to investigate whether sex and ancestry are associated with differences in PDAC molecular features. Our primary analytic approaches included: (i) global comparisons of tumour mutation burden, including single nucleotide variant (SNV) density and percent genome altered, (ii) gene-level comparisons of SNVs, copy number aberrations (CNAs) and mRNA abundance, and (iii) evaluation of *KRAS* variant frequencies.

### Data preprocessing

We assembled a cohort of 7,527 patients from five datasets, with SNV data available for all PDACs, and somatic CNAs and mRNA abundance data for 1,046 (13.9%) PDACs. Data was obtained from the ICGC data portal (TCGA, PACA-AU, PACA-CA; release date 2019-11-26), the GDC data portal (CPTAC; release date 2021-07-01) and cBioPortal (AACR GENIE; release date 2024-04-03) [2,3,6–8]. Sex was patient-reported. Patient-reported ancestry and previously determined genetic ancestry from the GDC data portal were used for TCGA and PACA-AU respectively. For PACA-CA and CPTAC, germline VCFs were processed using bcftools (v1.15.1), tabix (v1.20) and PLINK2 (v2.00). Germline VCFs for PACA-CA were obtained directly, while for CPTAC, they were generated from BAM files using GATK (v4.2.4.1, RRID:SCR_001876) through a standardized DNA pipeline [9,10]. Genetic ancestry was determined using ADMIXTURE (v1.3.0, RRID:SCR_001263) to classify patients as European (EUR), African (AFR), Admixed American (AMR) or Asian (ASN) resulting in 6,422 with available ancestry (**Supplemental Table 1**) [11]. For admixed individuals, ancestry was assigned to the population with the largest genetic contribution. Due to lower SNVs/Mbp in CPTAC compared to other datasets, SNVs were recalled using a standardized pipeline using Strelka2 and Mutect2 followed by annotation against GENCODE (RRID:SCR_014966) v45 with VEP[9,10] (https://github.com/uclahs-cds/metapipeline-DNA).

### Statistical analysis

Throughout we employed an established two-stage statistical analysis in which significant univariate results (FDR < 0.10) progressed to multivariable analyses adjusting for age, sex and ancestry [12–17]. Global measures of mutation density – SNV density and <u>p</u>ercent of the <u>g</u>enome with a copy number <u>a</u>berration (PGA) were analyzed using unpaired two-sided t-tests or Mann-Whitney U tests as appropriate for distributional assumptions. For per-gene analyses, infrequently detected genes were excluded to focus on candidate driver events (**Supplemental Table 2**). European ancestry had the largest sample size and was used as the reference for ancestry comparisons. Random effects meta-analysis was used to account for dataset differences for mRNA and CNA analyses using meta (v7.0.0) and metafor (v.4.6.0). All analysis were conducted in R (v4.2.0), using data.table (v1.15.4) and parallel (v4.2.0) for data manipulation and BPG (v7.0.3) for visualization.

### CNA analysis

CNA analyses excluded genes with CNAs in <5% of patients or on sex chromosomes. CNAs were discretized to gain, neutral or loss using dataset-specific thresholds.[2,3,6,7] PGA was used as a proxy for the copy number load for each tumour, and calculated as the number of base pairs affected by CNAs per patient divided by the number of base pairs covered. For gene-specific CNA analysis, CNA segments were converted to gene level events using biomaRt (v2.54.1, RRID:SCR_019214) and GenomicRanges (v1.50.2, RRID:SCR_017051).

### mRNA abundance analysis

mRNA analyses excluded genes with zero reads in >75% of samples, and log_2_ mRNA abundance was z-score transformed. Results were not strongly influenced by either this threshold or transformation.

### SNV analysis

SNV analyses excluded genes on sex chromosomes. SNV density was determined as SNVs per patient per Mbp sequenced (with targeted panel-specific adjustments for AACR GENIE). For gene-specific SNV proportion analyses, only genes mutated (missense or nonsense mutations) in ≥2% of patients were included. Ancestries with <63 patients were excluded: this sample size allowed at least 80% power to detect a Cohen’s h of 0.5 in a two-sample proportion test. Given the prognostic value of *KRAS* variants, *KRAS* variant frequency was evaluated in a subset analysis focused on variants present in ≥5% of patients [18].

## Results

### Somatic mutation density varies by ancestry

To evaluate broad genomic changes, we first assessed genomic instability. In many other cancer types, genomic instability is associated with ancestry or sex, even in small cohorts [16]. In this large PDAC cohort, point mutation density (SNVs/Mbp) varied by ancestry (**Fig. 1a**) but not by sex (**Fig. 1b**) in univariate analysis. After multivariable adjustment, this association remained significant for those of African ancestry (p = 0.0027) and became apparent for those of Asian ancestry (p = 0.008). On the average, there were ∼4,400 more SNVs in African *vs*. European tumours, and ∼2,100 fewer SNVs in Asian *vs*. European tumours. These relatively large differences are of interest given the increased incidence and mortality rates of PDAC in patients of African ancestry [19].

**Figure 1.**
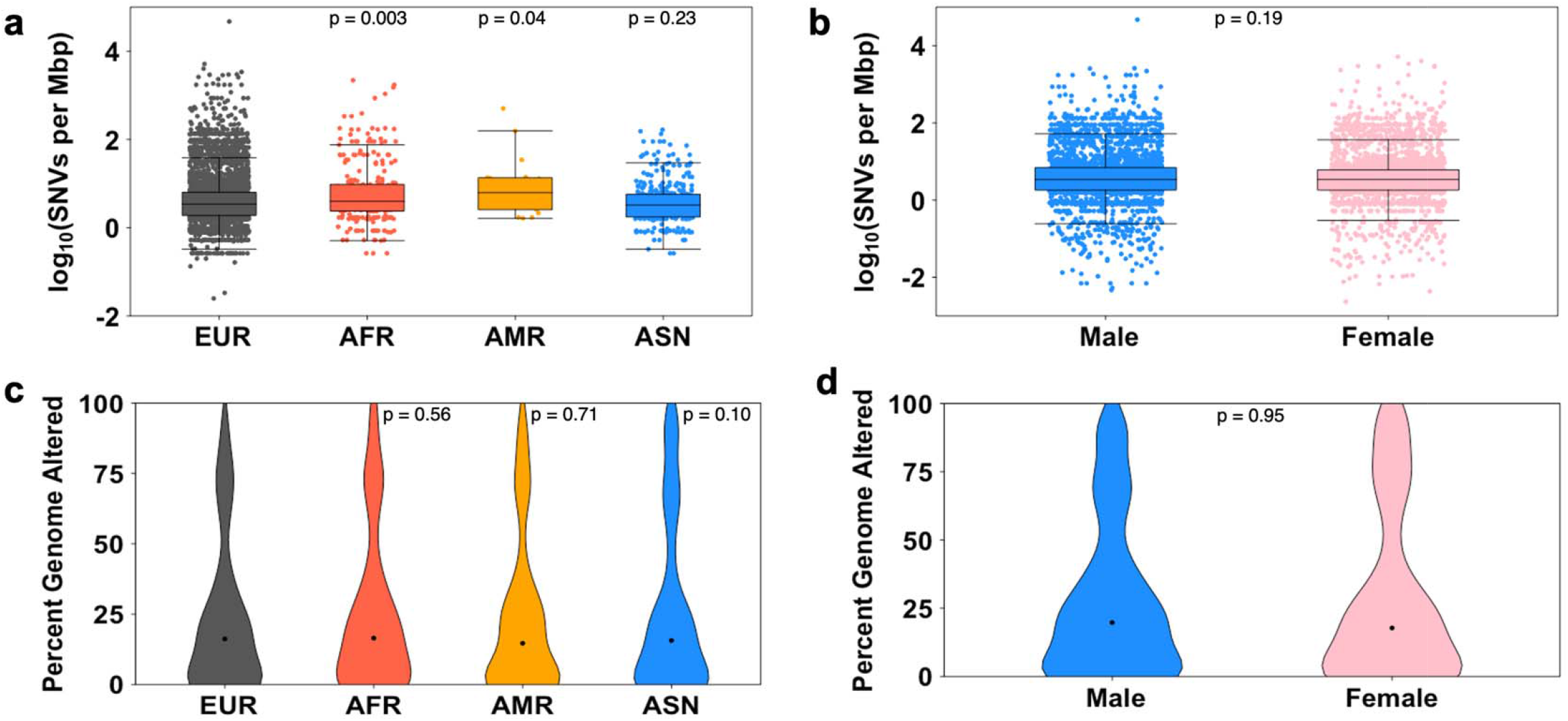
Mutation density and ploidy by sex and ancestry. SNV density differed across (a) ancestry but not (b) sex. No difference in percent genome altered was observed by (c) ancestry or (d) sex.

### Sex and ancestry have minimal influence on ploidy and mRNA profiles

Ploidy did not vary by either ancestry (**Fig. 1c**) or sex (**Fig. 1d**) even in univariate analyses. Similarly, bulk mRNA abundance profiles did not systematically vary by ancestry or sex. Of the 25,017 genes analyzed, none differed by ancestry, while 35 genes showed sex-associated differences in univariate analysis (**Fig. 2a**), of which only nine sex chromosome genes retained evidence of sexual dimorphism after multivariable adjustment (**Fig. 2b**). Intriguingly, some of these sex chromosome genes are relevant to PDAC biology. For example, reduced expression of *KDM5D* has been associated with adverse clinicopathological features and worse overall survival [20].

### Ancestry-associated differences in *GNAS* SNV frequency

We next focused on driver SNVs, which are very frequent in PDAC. Several specific genes showed modest effects including *CDKN2A* (**Fig. 3a**) and *FAT1* (**Fig. 3b**), with the most notable being a significant ancestry-bias in the oncogene *GNAS*. Driver SNVs in *GNAS* were detected in only 0.9% of patients of African ancestry, compared to 3.0% of those of European ancestry and fully 5.3% of those of Asian ancestry (**Fig. 3c**). This effect remained significant after both multiple-testing adjustment (q = 0.047) and multivariable adjustment for sex and age. *GNAS* is a potential alternative oncogenic driver in *KRAS* wild-type tumours [6,7] and is enriched in the classical subtype of PDAC, which has been associated with a more favorable clinical presentation [3,7].

**Figure 2.**
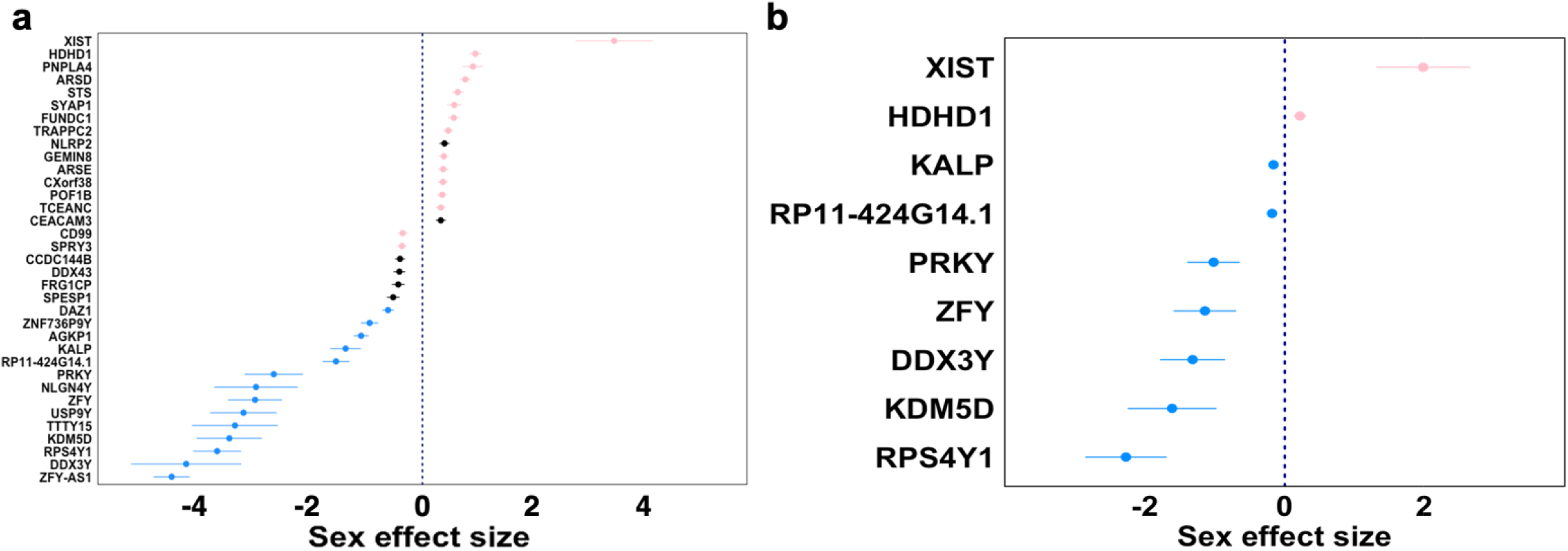
mRNA abundance by sex. (a) Thirty-five genes were significant in univariate analysis; (b) nine genes showed sex-specific differences in mRNA abundance after multivariable adjustment. Genes on the X and Y chromosome are depicted in pink and blue respectively. Negative values indicate genes with mRNA abundance that is lower in females compared to males, and vice versa.

**Figure 3.**
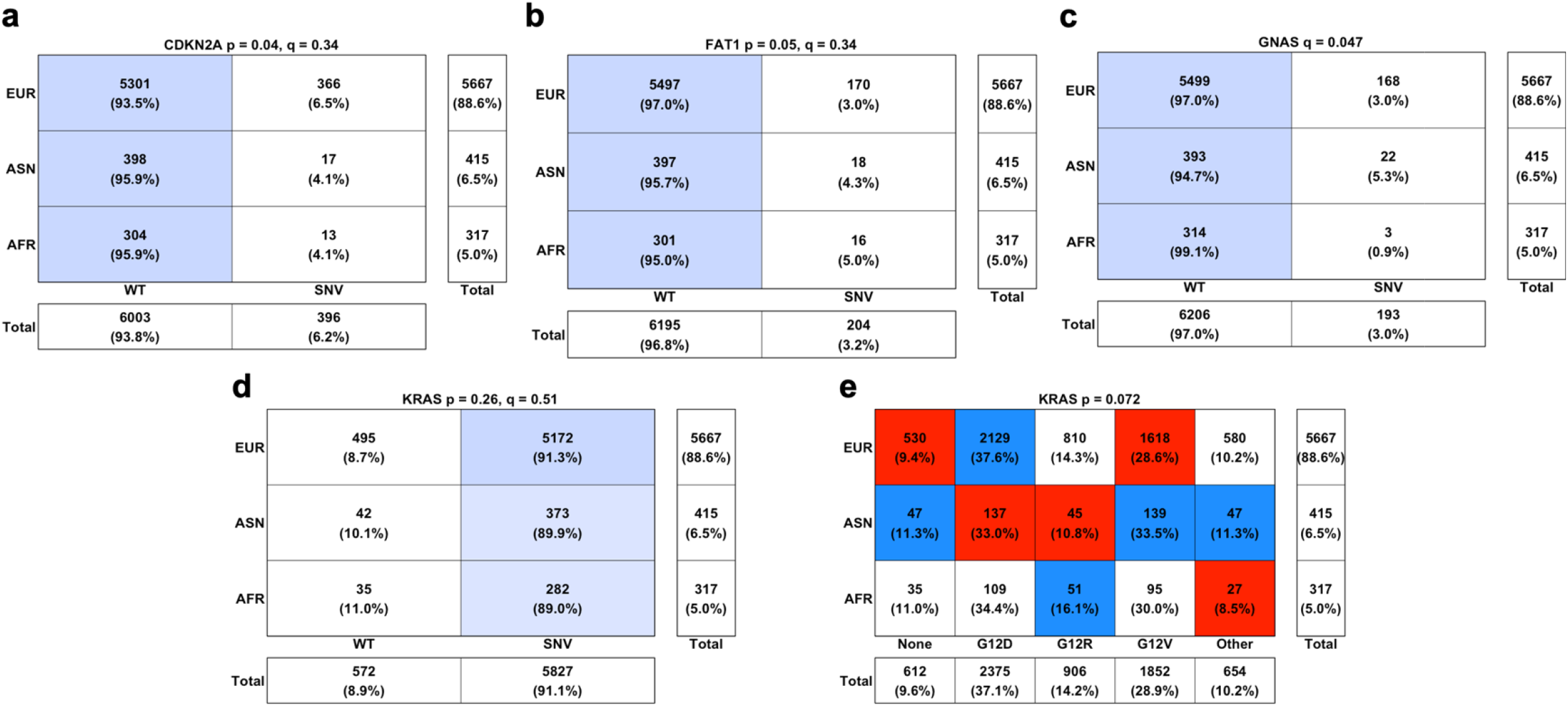
Driver SNV analysis by ancestry. Difference in SNV frequency across ancestry in (a) *CDKN2A*, (b) *FAT1* and (c) most significantly *GNAS*. There was no significant difference in *KRAS* SNV proportion across (d) ancestry. Color shading from white to blue indicates the relative contribution of each cell to the row total. Subset analysis revealed a trend towards *KRAS* codon-specific differences across (e) ancestry. For each column, red, white and blue indicates the ancestries with the lowest, intermediate and highest frequencies of the variant, respectively.

### Ancestry and sex influence *KRAS* variant distribution

This ancestry bias in *GNAS*, a proposed driver in *KRAS* wild type tumours, prompted us to next examine whether *KRAS* and its variants might also demonstrate ancestry or sex specific differences in PDAC. This is particularly relevant given the prognostic and therapeutic implications of certain *KRAS* variants [18]. The proportion of *KRAS*-negative tumours did not vary by either sex (q = 0.55; **Supplemental Fig. 1a**) or ancestry (**Fig. 3d**). By contrast, there was a strong trend towards specific *KRAS* codons being preferentially mutated in patients of specific ancestries (p = 0.072, **Fig. 3e**), and to a lesser extent sex (p = 0.10; **Supplemental Fig. 1b**). Poor-risk G12D variants were more common in individuals of European ancestry, while favorable risk G12R variants were more common in those of African ancestry. Most dramatically, individuals of Asian ancestry harboured 17% more G12V mutations than those of European ancestry. Poor-risk G12D variants were more common in males, while G12V and favorable risk G12R variants were more common in females.

## Discussion

In summary, we quantified in a large cohort the impact of sex and ancestry on the genomic landscape of PDAC. These are the two largest contributors to germline variation, and had relatively modest influences. No association with total copy number was identified, and effect-size associations with mRNA abundance, SNV density and specific driver mutations were generally few and modest compared to other cancer types. Nevertheless, the few associations observed are of great interest because of the increasing targetability of *KRAS* and other oncogenes. The most common *KRAS* variants in PDAC include G12D, G12V and G12R. G12R has been associated with significantly longer overall survival compared to G12D, suggesting that codon-specificity has clinical implications beyond targetability [18]. The recent FDA approval of daraxonrasib, a RAS-targeted therapy for metastatic pancreatic adenocarcinoma, highlights the growing clinical relevance of understanding variant-specific differences in *KRAS* mutant PDAC. Given that G12D is the most common *KRAS* variant in PDAC, the emergence of effective RAS-directed therapies further underscores the importance of understanding inter-tumoural heterogeneity in *KRAS* mutation acquisition and its potential clinical implications [21].

A key limitation of this study is the unequal representation of ancestry groups. Despite a large overall sample-size, patients of European ancestry were over-represented, limiting statistical power. Nevertheless, outside of *KRAS* allelic differences, analyses were sufficiently powered to detect large effect sizes. Tumour purity and stage (not examined in this work due to limited availability), may also influence results and warrants future study. Variations in sequencing coverage resulting from differences in specimen processing and technique across datasets are additional possible confounding factors. Moreover, it is important to consider that social determinants of health may contribute to disparities in both outcomes and molecular presentation.

## Conclusions

In summary, we demonstrated that ancestry influences the genomic heterogeneity of PDAC. These findings emphasize the importance of continued efforts to understandgermline-somatic interactions, particularly among underrepresented ancestral groups. Research focusing on less represented ancestral groups and sub-groups will be critical to enhance our understanding of PDAC heterogeneity, and could ultimately lead to more personalized treatment approaches and improved management of this highly lethal disease.

## Declarations

### Ethics approval and consent to participate

Not applicable

### Availability of data and materials

Requests for further information and resources should be directed to and will be fulfilled by the corresponding author. This study did not generate new materials. Scripts for data processing, analysis and visualization are available upon request.

### Authors’contributions

Project Initiation: C.Y.C., P.C.B. Bioinformatic Analyses: C.Y.C., C.Z., J.O.

Statistical Analyses: C.Y.C., J.A. Manuscript First Draft: C.Y.C. Supervised Research: P.C.B.

Manuscript Editing & Approval: All authors

## Acknowledgements

The authors thank all members of the Boutros Lab for their input and support.

## Abbreviations

(AFR): African
(AMR): Admixed American
(ASN): Asian
(CNA): Somatic copy number aberration
(EUR): European
(PDAC): Pancreatic ductal adenocarcinoma
(SNV): Single nucleotide variant

