## Supplemental Figures for "The impact of sex and ancestry on the molecular landscape of pancreatic adenocarcinoma"

**Supplemental Table 1: SNV dataset composition by sex and ancestry**

| | Male | Female | Other | Total |
|---|---|---|---|---|
| **EUR** | TCGA = 80<br>PACA-AU = 44<br>PACA-CA = 87<br>CPTAC = 61<br>AACR GENIE = 2736<br>**Subtotal = 3008** | TCGA = 65<br>PACA-AU = 41<br>PACA-CA = 75<br>CPTAC = 52<br>AACR GENIE = 2421<br>**Subtotal = 2654** | TCGA = 0<br>PACA-AU = 0<br>PACA-CA = 3<br>CPTAC = 0<br>AACR GENIE = 2<br>**Subtotal = 5** | 5667 |
| **AFR** | TCGA = 4<br>PACA-AU = 0<br>PACA-CA = 1<br>CPTAC = 1<br>AACR GENIE = 131<br>**Subtotal = 137** | TCGA = 3<br>PACA-AU = 0<br>PACA-CA = 3<br>CPTAC = 1<br>AACR GENIE = 173<br>**Subtotal = 180** | TCGA = 0<br>PACA-AU = 0<br>PACA-CA = 0<br>CPTAC = 0<br>AACR GENIE = 0<br>**Subtotal = 0** | 317 |
| **AMR** | TCGA = 3<br>PACA-AU = 1<br>PACA-CA = 1<br>CPTAC = 0<br>AACR GENIE = 8<br>**Subtotal = 13** | TCGA = 2<br>PACA-AU = 0<br>PACA-CA = 0<br>CPTAC = 0<br>AACR GENIE = 8<br>**Subtotal = 10** | TCGA = 0<br>PACA-AU = 0<br>PACA-CA = 0<br>CPTAC = 0<br>AACR GENIE = 0<br>**Subtotal = 0** | 23 |
| **ASN** | TCGA = 5<br>PACA-AU = 4<br>PACA-CA = 3<br>CPTAC = 12<br>AACR GENIE = 176<br>**Subtotal = 200** | TCGA = 6<br>PACA-AU = 4<br>PACA-CA = 6<br>CPTAC = 13<br>AACR GENIE = 186<br>**Subtotal = 215** | TCGA = 0<br>PACA-AU = 0<br>PACA-CA = 0<br>CPTAC = 0<br>AACR GENIE = 0<br>**Subtotal = 0** | 415 |
| **Total** | 3358 | 3059 | 5 | 6422 |

**Supplemental Table 2: Summary of methods**

| Analysis | Model | Univariate threshold | Multivariable threshold | Exclusion criteria |
|---|---|---|---|---|
| PGA by sex | Unpaired t-test | 0.10 | 0.05 | Genes with CNA in < 5% of patients or on sex chromosomes |
| SNV density by sex | Unpaired t-test | 0.10 | 0.05 | Genes on sex chromosomes |
| PGA by ancestry | Unpaired Mann-Whitney U test | 0.10 | 0.05 | Genes with CNA in < 5% of patients or on sex chromosomes |
| SNV density by ancestry | Unpaired Mann-Whitney U test | 0.10 | 0.05 | Genes on sex chromosomes |
| mRNA abundance | Random effects meta-analysis (Hedges' g by sex, linear regression by ancestry) | 0.10 | 0.05 | Genes with zero reads in more than 75% of samples |
| Gene-specific CNA gains or losses | Random effects meta-analysis (relative risk by sex, logistic regression by ancestry) | 0.10 | 0.05 | Genes with CNA in < 5% of patients or on sex chromosomes |
| Gene-specific SNV proportion analysis by sex | Two-sample proportion test | 0.10 | 0.05 | Silent mutations, genes mutated in < 2% of patients or on sex chromosomes |
| Gene-specific SNV proportion analysis by ancestry | Fisher's Exact test | 0.10 | 0.05 | Silent mutations, genes mutated in < 2% of patients or on sex chromosomes<br>Ancestries with fewer than 63 patients |
| *KRAS* variant proportion analysis by sex | Fisher's Exact test | 0.10 | 0.05 | Silent mutations, variants present in < 5% of patients |
| *KRAS* variant proportion analysis by ancestry | Fisher's Exact test | 0.10 | 0.05 | Silent mutations, variants present in < 5% of patients<br>Ancestries with fewer than 63 patients |

### Supplemental Figure 1: (a) *KRAS* SNV frequency and (b) *KRAS* variant SNV frequency by sex

**a**

KRAS OR = 1.08, p = 0.28, q = 0.55

| | Male | Female | Total |
|---|---|---|---|
| WT | 370 (54.3%) | 312 (45.7%) | 682 (9.1%) |
| SNV | 3590 (52.4%) | 3255 (47.6%) | 6845 (90.9%) |
| Total | 3960 (52.6%) | 3567 (47.4%) | |

**b**

KRAS p = 0.103

| | None | G12D | G12R | G12V | Other | Total |
|---|---|---|---|---|---|---|
| Males | 345 (9.0%) | 1503 (39.0%) | 520 (13.5%) | 1097 (28.5%) | 387 (10.0%) | 3852 (52.6%) |
| Females | 280 (8.1%) | 1289 (37.2%) | 523 (15.1%) | 1034 (29.8%) | 341 (9.8%) | 3467 (47.4%) |
| Total | 625 (8.5%) | 2792 (38.1%) | 1043 (14.3%) | 2131 (29.1%) | 728 (9.9%) | |
